# Back to feedback: aberrant sensorimotor control in music performance under pressure

**DOI:** 10.1101/2020.05.16.100040

**Authors:** Shinichi Furuya, Reiko Ishimaru, Takanori Oku, Noriko Nagata

## Abstract

Precisely timed production of dexterous actions is often destabilized in anxiogenic situations. Previous studies demonstrated that cognitive functions such as attention and working memory as well as autonomic nervous functions are susceptible to induced anxiety in skillful performance while playing sports or musical instruments. However, it is not known whether the degradation of motor functions, sensory perception, or sensorimotor control underlies such a compromise of skillful performance due to psychophysiological distress. Here, we addressed this issue through a series of behavioral experiments, which provided no evidence supporting for detrimental effects of the stress on the perceptual accuracy and precision of the finger movements in pianists. By contrast, after transiently delaying the timing of tone production while playing the piano, the local tempo was abnormally disrupted only under pressure. The results suggest that psychological stress degraded the temporal stability of movement control due to an abnormal increase in sensory feedback gain but not temporal perception or motor precision. A learning experiment further demonstrated that the temporal instability of auditory-motor control under pressure was alleviated after practicing piano while ignoring delayed auditory feedback but not after practicing while compensating for the delayed feedback. Together, these findings suggest an abnormal transition from feedforward to feedback control in expert piano performance in anxiogenic situations, which can be mitigated through specialized sensorimotor training that involves piano practice while volitionally ignoring the artificially delayed provision of auditory feedback.

## INTRODUCTION

Skillful behaviors are readily spoiled in anxiogenic situations such as the Olympic games and music competitions. Suboptimal performance of skillful actions due to psychological stress and fear under pressure is a common problem across trained individuals, such as athletes, musicians, and surgeons, and sometimes affects the future career prospects of these individuals. State anxiety typically compromises spatiotemporal control of dexterous movements (1-6). For example, stuttering in speech is exaggerated under pressure (7), which indicates the detrimental effects of psychological stress on precisely timed execution of motor sequences. Similarly, psychological stress triggers rhythmic distortion of sequential finger movements in musical performance (8-10). These deleterious effects of psychological stress on motor precision have been extensively argued in terms of their relation to abnormal attention under pressure (1, 5, 11-14). For example, skilled baseball players were more aware of the direction of bat motion in a higher pressure condition, and it was speculated that the exaggerated attention to the well-learned processes of skill execution led to abnormal modulation of skill execution and thereby increased the movement variability under pressure (5). In contrast to cognitive malfunctions such as abnormal attention, it has not been elucidated what malfunctions in sensory-motor processes underlie the degradation of precision of skillful motor actions due to psychological stress. Understanding this is essential to shed light on neuropsychological mechanisms subserving the robustness and flexibility of sensorimotor skills in various environments.

At least three malfunctions of the sensorimotor system are candidates that may compromise timing control of motor sequences due to psychophysiological distress. The first candidate is abnormal temporal perception due to psychological stress. It has been reported that fear and anxiety distort time perception, such as interval timing and simultaneity (15-17). For example, a recent study showed that experimentally inducing anxiety resulted in underestimating the duration of temporal intervals (17). The second candidate is abnormal motor functions under pressure. It has been reported that induced stress elevated finger force fluctuation during a precision grip (18, 19), which suggests the degradation of motor functions in stressful situations. However, most of these previous studies targeted individuals without any history of extensive training, which questions the time perception and motor functions of experts under pressure.

The third candidate is sensory feedback control of movements. A key element of skillful behaviors is the robustness of skills against external perturbation. The balance between the robustness and flexibility of sensorimotor skills dynamically shifts over a course of learning (20-23). At the early stage of skill acquisition, a large gain of sensory feedback enables fast skill improvements with training. Through extensive training, such a skill reaches its plateau and is stabilized. The stabilized skill is minimally susceptible to sensory perturbations due to feedforward control, which has been demonstrated in trained individuals (24). For example, singers but not nonmusicians can keep singing accurately even by artificially shifting the pitch of the voice (25) or by blocking somatosensory afferent information via vocal-fold anesthesia (26). Similarly, during piano playing, delayed tone production perturbs the subsequent keystroke motions to a lesser extent in expert pianists than in nonmusicians (27). Accordingly, it is possible that psychological pressure destabilizes experts’ sensorimotor control, which mediates abnormal increases in the timing fluctuation of skilled motor actions under pressure (9).

Here, we addressed the effects of psychological stress on time perception, motor functions, and sensorimotor control of expert pianists. We focused on musicians because many musicians suffer from state anxiety when performing on the stage, so-called performance anxiety (28, 29), which occurs in up to 90% of musicians. The stability of sensorimotor control was assessed by the well-established experimental paradigm using the altered sensory feedback, which provided transient sensory perturbation during piano playing (30). By using sensory perturbation, we also tested whether sensorimotor training can alleviate the detrimental effects of psychological stress on sensory-motor skills.

## MATERIALS & METHODS

### Participants

In the present study, 11 (7 females, 25.4 ± 4.9 yrs old) and 30 (23 females, 24.1 ± 6.0 yrs old) right-handed expert pianists with no history of neurological disorders participated in experiments 1 and 2, respectively. All of the pianists majored in piano performance in a musical conservatory and/or had extensive and continuous private piano training under the supervision of a professional pianist/piano professor. The age at which the participants had started to play the piano was before 8 years old. In accordance with the Declaration of Helsinki, the experimental procedures were explained to all participants. Informed consent was obtained from all participants prior to participating in the experiment, and the experimental protocol was approved by the ethics committee of Sophia University.

### Experimental Design

The present study consists of 2 experiments under both low-pressure (LP) and high-pressure (HP) conditions. To circumvent confounding effects of familiarization with the experimental situation and stress induction, participants were different across the experiments.

Experiment 1 was designed to test whether psychological stress influences the perception of the duration of time interval based on auditory stimuli, the agility and accuracy of movements, and sensory feedback control of movements in piano performance. In total, 3 tests were carried out, which assessed the perceptual threshold for discrmination of the auditory time interval (“*perception test*”), finger motor agility and accuracy (“*motor test*”), and auditory feedback control of movements (“*sensorimotor test*”) in the LP and HP conditions.

The perception test provided 2 sets of auditory stimuli, each of which consisted of 2 successive tones. The first “reference” stimulus was 2 successive tones with the inter-tone duration of 129.3 msec (i.e. a tempo of 116 beats per minute (bpm), which is same as one in the sensorimotor task), whereas the second stimulus was 2 successive tones with the inter-tone duration specified according to the staircase method. After providing the 2 stimuli, participants were asked to answer whether the inter-tone duration of the second stimulus was shorter than that of the first stimulus. Here, the inter-tone duration of the second stimulus was determined by a three-down one-up staircase method. At the first trial, the inter-tone duration was set to 123.0 ms. At the next trial, the inter-tone duration was decreased or increased by 2 ms, according to whether correct answers were repeated 3 times or an answer was incorrect once, respectively. The test was terminated when 4 peaks and 4 troughs were measured, and the average of the last 2 peaks and 2 troughs of the inter-tone duration was defined as the perceptual threshold of the auditory time-interval discrimination. In the present perceptual test, we employed this method rather than estimating the psychometric function because the former was quicker in the threshold detection and the pilot experiment found that the pressure effect tended to drop over time.

The motor test asked the participants first to perform repetitive strikes of a piano key with each of five digits of the right hand as fast and accurately as possible for 5 seconds (“the fastest single finger tapping”) (31) and second to play the first 8 tones of the piece of Op. 10 No. 8, composed by Frédéric François Chopin, on the piano as fast and accurately as possible (“the fastest piano performance”). Here, the second motor task consisted of tones of A5, G5, F5, C5, A4, G4, F4, and C4, which were played with the fingering of the ring, middle, index, thumb, ring, middle, index, and thumb, respectively. During the performance of these motor tests, the piano tones were deprived (i.e., muted) so that auditory feedback could not affect movement execution. These motor tests evaluated both the speed and the timing precision of the sequential keystrokes with a single finger and with multiple fingers by computing the mean and the standard deviation of the inter-keystroke interval across the strikes within a trial. Each task was repeated twice.

The sensorimotor test asked the participants to play a short melody extracted from Op. 10 No. 8, composed by Frédéric François Chopin, with only the right hand (the first 14 bars) on an acoustic piano with MIDI sensors (YUS1SH, Yamaha, Co.) at a tempo of 116 bpm (interkeystroke interval = 129.3 msec) and at the loudness of *mezzo-forte*. This piece was chosen first because it is technically challenging enough to elicit anxiety for many pianists (note that this piece has been played at many piano competitions and during the entrance exams for many musical conservatories) and second because it requires high timing precision of the keystrokes during fast tempo performance with little effects of emotional or aesthetic expression (9). In addition, we previously confirmed disruption of performance of the present task by the induced psychological pressure (9). The target melody was to be played with legato touch, meaning that a key was not released until the next key was depressed. On the day of the experiment, participants were initially asked to become familiar with the piano by practicing for 10 min, without using a metronome, prior to the experiment. After the familiarization session, all participants were able to play the target melody with the predetermined fingering at the target tempo without pitch errors. During the sensorimotor test, prior to the task performance, 16 metronome tones were provided to a cue of the tempo (i.e., 116 bpm), and then the pianists were asked to play without the metronome sounds. While playing the target melody, the timing of tone production was artificially delayed by 80 ms over 4 successive strikes (i.e., delayed auditory feedback) as auditory perturbation (27, 30, 32), which occurred 3 times within the melody. The notes with delayed tone production were not told to the pianists in advance and were randomized across the pianists so that each pianist could not predict which keystrokes elicited delayed tone production. It is obvious that such an artificially delayed tone production does not occur in a real piano performance on stage, but this empirical technique provides a unique opportunity of evaluating stability of the movements through assessing the relationship between the input (i.e. perturbation) and output (i.e. reaction) (30).

For experiment 1, the order of the perception, motor, and sensorimotor tests was randomized across the participants. In the LP condition, the experimenter sat behind each pianist and out of sight. In the HP session, the stress induction was performed in a manner used in our previous study (8). Another experimenter appeared and adopted a strict and unfriendly behavior; this experimenter stood next to the participant and monitored and evaluated the performance. In addition, the performance was videotaped by a camera mounted in front of the performer. Prior to initiating the HP session, the experimenter told the participants that he/she would evaluate the performance. This experimental design was validated and adopted in our previous studies (8, 9). In addition, in the present study, the participants were asked in each of the LP and HP conditions about to what extent they perceived the stress by using the 7-point Likert scale (a larger value indicates higher stress). The results of the subjective rating were 2.0 ± 1.0 and 5.7 ± 0.8 points in the LP and HP conditions, respectively (p = 2.06 × 10^−6^), which confirmed a significant difference in the stress level between the present experimental conditions. The order between the LP and HP conditions was randomized across the participants. Between the conditions, the participants took rest for about ten minutes until the heart rate returned to the baseline level.

Experiment 2 was designed to assess the effects of short-term piano practicing with the provision of transiently delayed auditory feedback on the disruption of piano performance following auditory perturbation. The experiment consisted of 4 successive sessions in the following order: playing in the LP condition (LP_pre_), intervention, playing in the LP condition (LP_post_), and playing in the HP condition (HP). Thirty pianists were randomly assigned to 3 groups undergoing different interventions. At the intervention session, they played the first 8 notes of the target melody over 200 trials. The first control group was asked to play the 8 notes without any artificial delay in the timing of the provision of auditory feedback (“normal” group). The second group was instructed to play while ignoring the 80-ms delayed tone production, which occurred at each of the last 4 notes (“ignore” group). The third group was instructed to play while compensating for the delayed tone production at each of the last 4 notes so that the tones could sound with consistent intertone intervals across the 8 notes (“compensate” group). This means that the pianists in this group had to strike the keys 80 ms prior to the timing at which the tone should be elicited at the latter 4 strikes. Prior to each trial, 4 metronome tones were provided as a cue for the target tempo, and then the participants began playing without the metronome cue. The intervention and its instruction was designed based on previous studies in which participants were instructed to either ignore or compensate for the altered auditory feedback during movements (25, 33, 34).

### Data Acquisition

We recorded MIDI data from the piano by using a custom-made script written in JAVA (27). This script allows for artificially delaying the timing of tone production during piano playing by waiting for a predetermined duration until the MIDI signal is elicited to the piano. Based on the MIDI data collected by this script during the experiments, we derived the time at which each key was depressed and released. Based on this information, the interkeystroke interval was computed (note that this is a difference in timing between two successive keypresses, but not a difference in tone onset between two successive tones). The electrocardiogram data of each pianist was also recorded during the task performance in the experiments (Mio Slice, Mio Global Inc.) and data were analyzed in an offline manner. The electrocardiogram data were then expressed in bpm (beat per minute) for each R-R-interval. In experiment 1 only, the heart rate recordings from 3 pianists were contaminated by unpredictable noises, resulting in missing data; therefore, the data from these pianists were not used for the subsequent analyses.

### Data Analysis

Using information on the interkeystroke interval during piano playing, we defined the timing error as the temporal gap of the interkeystroke intervals between the measured and ideal presses. Here, the interkeystroke interval of the ideal presses was defined based on the tempo provided by the metronome. The timing error for each of the interkeystroke intervals was then averaged across the strikes for each participant under each of the HP and LP conditions. For the perception test of experiment 1, the perceptual threshold was computed based on the aforementioned manner (i.e. a staircase method). For the motor test of experiment 1, the inter-keystroke interval was first computed as a difference in the timing of the keypress between two successive keystrokes, and secondly averaged across the keypresses within a trial. The derived mean inter-keystroke interval within a trial was averaged between the two trials, and the reciprocal of the averaged inter-keystroke interval was defined as the tap rate in Hz (i.e. taps per second).

### Statistics

For datasets of the perception test and motor test (the fastest piano performance) in experiment 1, a paired t-test was performed for a comparison between the HP and LP conditions (p = 0.05). If the dataset did not follow the normal distribution based on the Kolmogorov-Smirnov test, a nonparametric Wilcoxon test was performed. For the heart rate in experiment 1, a two-way repeated-measures ANOVA was performed with task (3 levels: perception, motor, sensorimotor tests) and condition (2 levels: LP and HP) as within-subject independent variables. For the fastest finger tapping at the motor test in experiment 1, a two-way repeated-measures ANOVA was performed with digit (5 levels: thumb, index, middle, ring, and little fingers) and condition (2 levels: LP and HP) as within-subject independent variables. For the timing error at the sensorimotor test in experiment 1, a two-way repeated-measures ANOVA was performed with strike (4 levels: the first, second, third, and fourth strikes following provision of the delayed tone production) and condition (2 levels: LP and HP) as within-subject independent variables. With respect to experiment 2, a two-way mixed-design ANOVA was performed with group (3 levels: normal, ignore, and compensate) and strike (4 levels: the first, second, third, and fourth strikes following the provision of delayed tone production) as independent variables. Mauchly’s test was used to test for sphericity prior to performing each ANOVA, and for nonspherical data, the Greenhouse-Geisser correction was performed. Post hoc tests with correction for multiple comparisons (35) were performed in the case of significance. These statistical analyses were performed with R statistical software (ver. 3.2.3.).

## RESULTS

### Effects of psychological stress on perceptual, motor, and sensorimotor functions

Figure 1 illustrates the summarized results of experiment 1 that assessed the effects of psychological stress on heart rate and the perception of the time interval (i.e. duration perception), finger motor dexterity (i.e., motor function), and feedback control of movements based on sensory information (i.e., sensorimotor control) in the pianists.

**Figure 1.**
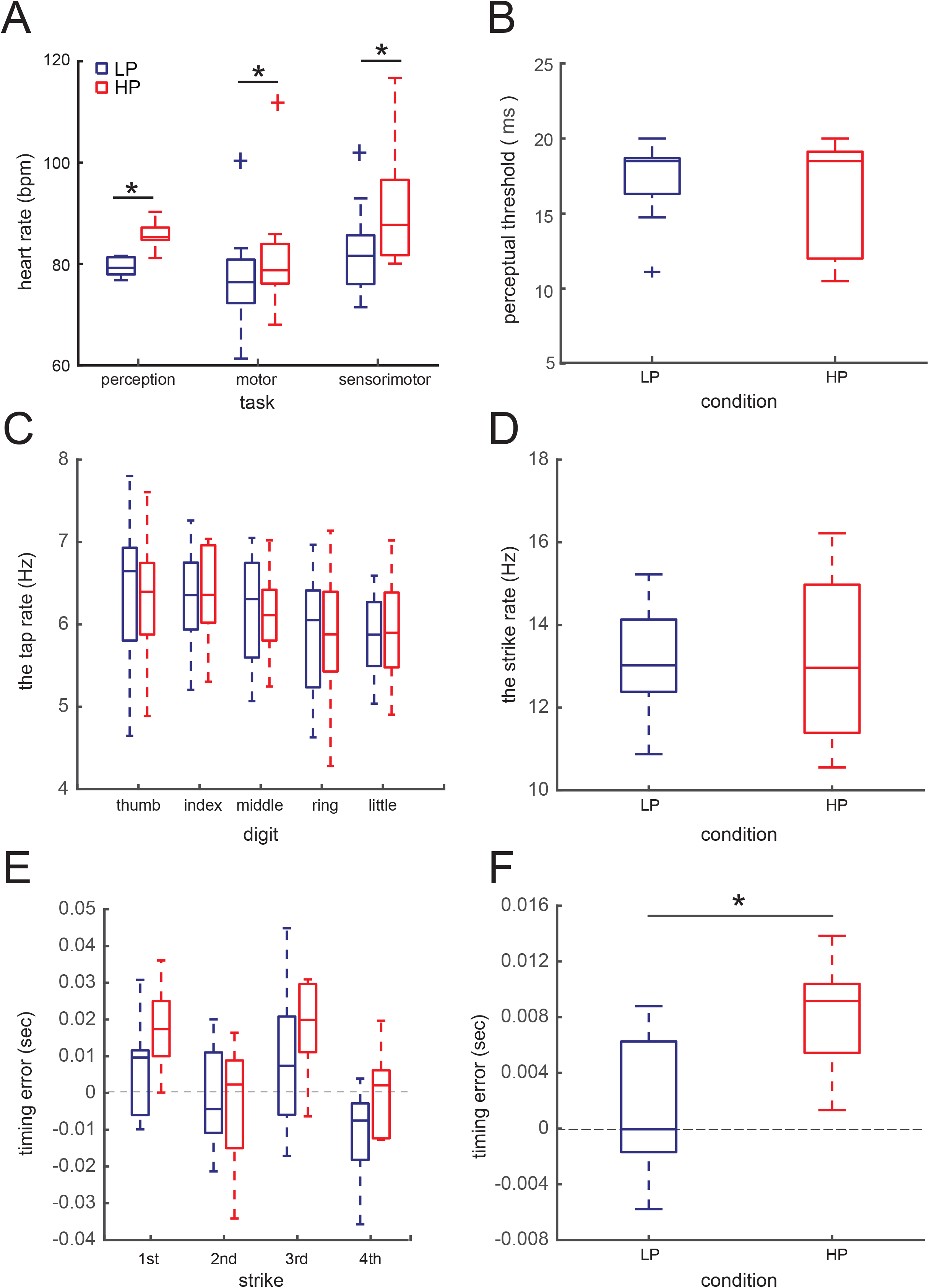
Results of the experiment 1. (A) Group means of the average heart rate during the perception, motor, and sensorimotor control tests; (B) the perceptual threshold of time duration between two tones; (C) the tap rate during the fastest tapping by each of five digits (i.e. taps per sec); (D) the strike rate by the fingers while playing the short melody excerpt as fast as possible without auditory feedback being provided (i.e. strikes per sec); (E) the timing error of the keypresses over 4 strikes following the provision of transiently delayed tone production, and (F) their mean across four strikes in the low-pressure (LP) and high-pressure (HP) conditions. *: p < 0.05. The blue and red boxplots indicate the LP and HP conditions, respectively.

Figure 1A displays the group means of the average heart rate during each of the perception, motor, and sensorimotor control tasks in the pianists in the HP and LP conditions. During each of the three tasks, the heart rate was higher for the HP condition than for the LP condition. A two-way repeated-measures ANOVA demonstrated a main effect of condition (2 levels: HP and LP) on heart rate (F(1, 7) = 22.71, p = 0.002, η^2^ = 0.116), but neither a main effect of task (3 levels: task 1-3; F(2, 14) = 2.66, p = 0.11, η^2^ = 0.121) nor an interaction effect (task × condition: F(2, 14) = 2.47, p = 0.12, η^2^ = 0.008) was evident. A lack of the interaction effect indicates that the pressure effect on the heart rate did not differ between the three different tasks.

Figure 1B illustrates the results of the psychophysics experiment that evaluated the threshold of the auditory perception of the duration of the time interval between two successive tones in pianists in the HP and LP conditions (i.e., *the perception test*). A paired t-test yielded no significant difference between the conditions (t(10) = 1.163, p = 0.136). This indicates no effects of psychological stress on the auditory perception of the time interval between the two tones.

Figure 1C displays the group means of the maximum tapping rate (i.e. a reciprocal of the mean inter-tap interval) by each of the five digits in pianists during the HP and LP conditions (i.e. *the motor test*). Here, the tap rate was computed as the reciprocal of the mean inter-keystroke interval between the successive keypresses within a trial. A two-way repeated-measure ANOVA yielded neither an interaction effect between finger and condition (F(4, 40) = 0.154, p = 0.96, η^2^ = 9.60 × 10^−5^) nor a main effect of condition (F(1, 10) = 3.16 × 10^−4^, p = 0.99, η^2^ = 5.57 × 10^−7^) but a significant main effect of finger (F(4, 40) = 6.27, p = 5.14 × 10^−4^, η^2^ = 0.10). This indicates that finger tapping ability did not differ irrespective of the pressure level. Similarly, with respect to the variability of the tap rate within a trial during the fastest finger tapping, neither an interaction effect between finger and condition (F(4, 40) = 0. 74, p = 0.570, η = 0.015) nor a condition effect (F(1, 10) = 0.40, p = 0.539, η = 0.002) were observed. This indicates no effect of psychological stress on the timing precision of the fastest finger tapping movements.

Figure 1D displays the maximum keystroke rate (i.e. a reciprocal of the mean inter-keystroke interval) while playing an excerpt of a melody that consists of 8 successive tones as fast as possible without auditory feedback being provided (i.e., *the motor test*). A paired t-test yielded no significant difference between the conditions (t(10) = 0.05, p = 0.962). Similarly, with respect to the variability of the keystroke rate within a trial while playing this melody as fast as possible, a paired t-test identified no significant difference between the conditions (t(10) = -0.37, p = 0.721). These results indicate no effects of psychological stress on both the speed and the accuracy of playing a short melody.

Figure 1E illustrates the timing error of the piano keypresses following the provision of the artificially delayed tone production in the pianists during the HP and LP conditions (i.e., *the sensorimotor test*). A positive value indicates transient slow-down of the local tempo following the perturbation. A two-way repeated-measure ANOVA yielded significant main effects of both condition (F(1, 10) = 19.46, p = 0.01, η^2^ = 0.073) and strike (F(3, 30) = 6.01, p = 0.002, η^2^ = 0.27) but no interaction effect of condition and strike (F(3, 30) = 2.11, p = 0.120, η^2^ = 0.06). To describe the main effect of the condition, the average value across the 4 strikes was computed for each of the HP and LP conditions (Fig. 1F). The effect of psychological stress on the timing error of the keypresses was characterized by the larger timing error in the HP condition compared with the LP condition. These results indicate that pianists slowed down the local tempo of piano playing following the transiently delayed tone production to a larger extent in a higher pressure condition.

### Effects of sensorimotor training

Experiment 2 tested whether practicing with the provision of delayed auditory feedback would influence the rhythmic distortion of the keypresses following the perturbation in piano performance under pressure. Figure 2 illustrates differences in the effects of psychological stress on motor responses to transiently delayed tone production among the 3 groups with different interventions (i.e. the normal, ignore, and compensate groups). Here, the value at each strike indicates a difference in the timing error of the keystrokes between the HP condition and the LP_pre_ condition, which was computed to minimize the confounding effects of possible individual differences in the interkeystroke interval across the groups. A two-way mixed-design ANOVA demonstrated main effects of both group (F(2, 27) = 5.85, p = 0.008, η^2^ = 0.07) and strike (F(3, 81) = 7.88, p = 1.12 × 10^−4^, η^2^ = 0.20) but no interaction effect between them (F(6, 81) = 1.17, p = 0.33, η^2^ = 0.07). A groupwise comparison with corrections for multiple comparisons identified a significant group difference only between the ignore and normal groups (inset of Fig. 2). Furthermore, there was no significant group effect on the timing error of the keystrokes in either the LP_pre_ condition (F(2, 27) = 1.26, p = 0.30, η^2^ = 0.02) or the LP_post_ condition (F(2, 27) = 2.90, p = 0.07, η^2^ = 0.05). Together, these findings indicate that the slow-down of the local tempo following delayed tone production under pressure was reduced after short-term piano practice while ignoring the delayed tone production.

**Figure 2.**
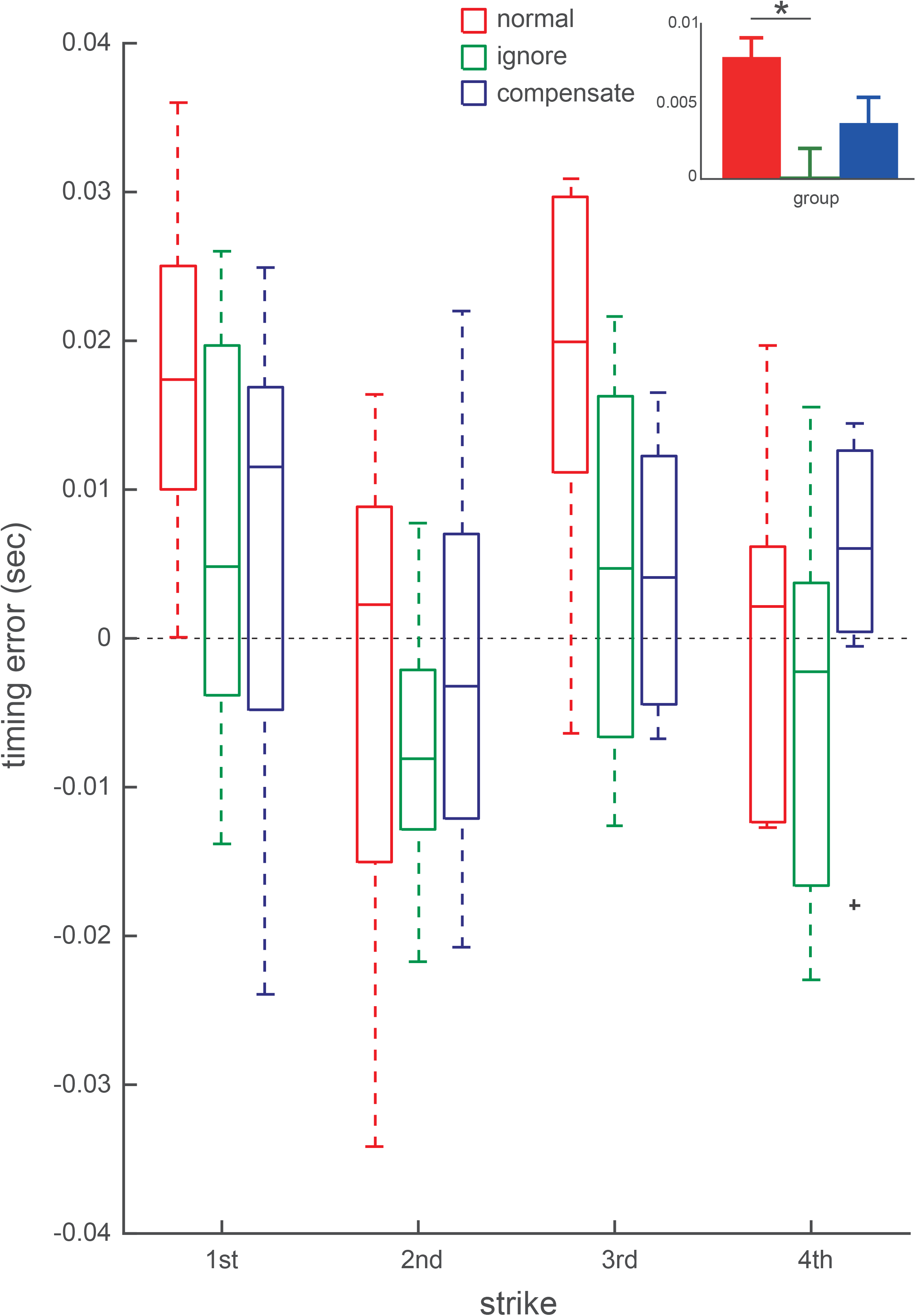
Results of the experiment 2. Group means of differences in the timing error of the four successive keypresses following the provision of transiently delayed tone production between the high-pressure (HP) condition and low-pressure condition before running the intervention (LP_pre_) for the normal (red), ignore (green), and compensate (blue) groups. x-axis: four successive strikes immediately after the transiently delayed tone production. The inset indicates the group means of the timing error across the four strikes for each of the normal, ignore, and compensate groups, which was depicted based on a significant group effect but not an interaction effect between group and strike (an error bar indicates one standard error). *: p < 0.05. The value 0 indicates no difference in the keypress timing error between the HP and LP_pre_ conditions.

To compare the heart rate between the groups, a two-way mixed-design ANOVA was performed by using the group (3 groups) and conditions (LP_pre_ and HP) as independent variables. Neither the interaction (F(2, 30) = 0.01, p = 0.986, η^2^ = 0.001) nor a group effect (F(2, 30) = 0.19, p = 0.829, η^2^ = 0.01) was significant, which indicated that the training with the delayed tone production did not affect the pressure effect on the heart rate. In addition, although the experiment 2 did not randomize the order of the LP and HP conditions because of the training experiment paradigm, the heart rate in the HP condition did not differ between the experiment 1 (the sensorimotor test) and experiment 2 (the normal group) (p = 0.097), which did not provide physiological evidence supporting for an order effect of the pressure effect.

## DISCUSSION

In the present study, we first tested whether psychological stress yields perceptual, motor, or sensorimotor malfunctions in pianists. The results demonstrated larger rhythmic disruption of the sequential finger movements following transient perturbation of auditory feedback at the higher pressure condition, which provided evidence supporting the abnormality in sensory feedback control of movements under pressure. However, neither auditory perception of time interval nor finger motor precision was affected by the induced psychological stress in the present experiments. These findings indicate that an increase in the timing error of sequential movements under pressure is associated with an abnormal elevation in the sensitivity of motor actions to perceived error (i.e., an elevated feedback gain in sensorimotor control), but not with abnormalities of the perception of the time interval or of fine motor control. It is unlikely that this difference in the pressure effect between the tasks was due to a difference in the task difficulty, due to a lack of any interaction effect of the condition and task on the heart rate (i.e. the pressure effect on the heart rate was not different between the three tests). In a nonstressful musical performance, the feedback gain of sensorimotor control is usually low for pianists but not for musically untrained individuals, as exemplified by a transient increase in rhythmic error of movements following a transient delay in the timing of auditory feedback during piano playing only in the latter individuals (27, 30). Therefore, these findings suggest the de-expertise of sensorimotor control of skilled pianists under pressure, which fits with the idea of choking, describing the failure of skill execution under pressure (36). In the subsequent experiment, we also tested whether sensorimotor training normalizes the instability of sensorimotor control under pressure. The results of the intervention experiment demonstrated that practicing the piano while ignoring artificially delayed tone production resulted in the amelioration of the unstable sensorimotor control under pressure. Intriguingly, this intervention effect was not observed when playing without stress, which indicates specific training effects on sensorimotor control during playing under stress. Furthermore, such an alleviation of the stress effect was not observed after practicing with compensation for delayed tone production and after practicing without the provision of delayed auditory feedback. Taken together, our results indicated that state anxiety abnormally augmented reliance on sensory feedback in the fine control of sequential finger movements during piano playing, which can be restored specifically by ignoring auditory perturbation.

A question is why the pianists transiently slowed down the tempo following the perturbed auditory feedback only in the anxiogenic situation. One may consider that the feedback gain was elevated due to an increased uncertainty of the perception under pressure. However, our results do not support this idea due to a lack of changes in auditory perception of the time interval under pressure. Another possibility is that the pianists became sensitive to the exogenous disturbance of auditory feedback in piano playing only under pressure, which had the pianists fail to ignore the disturbance. However, this is also not likely because of no difference in the amount of elevation in the heart rate due to the pressure between the present experiment with auditory perturbation (7.2 ± 2.2 bpm) and our previous experiment that used the same task without auditory perturbation (10.8 ± 6.8 bpm) (9) (p=0.129). One plausible explanation is that the transient slow-down of the local tempo following the perturbation reflected attempts to lower accuracy demands on the performance by leveraging a tradeoff between the speed and the accuracy of movements (37). This idea is compatible with elevated coactivation between the finger flexor and extensor muscles under pressure, which is considered a coping strategy for accuracy demand (8, 10). A recent study also proposed a role of muscular coactivation in feedback control of movements (38). The shift from feedforward to feedback control in the anxiogenic situation can affect reduced movement individuation between the fingers under pressure (9) because the elevation of the muscular coactivation increases the stiffness of the finger muscles connected with multiple fingers and thereby increases biomechanical constraints on the individuated finger movements (39). Importantly, the reduced movement individuation between the fingers was associated with an increase in the timing error of the piano performance under pressure (9), which suggests that elevating feedback gain in the stressful condition played no beneficial role in accurate musical performance. In addition, the transient slow-down of the local tempo after delayed tone production under pressure was also unlikely to contribute to accurate tempo control because the maintenance of the global tempo after delayed tone production requires a transient speed-up of the local tempo to compensate for the transient delay (30). Taken together, the increased reliance on feedback control under pressure is considered not as a strategy compensating for the external sensory perturbation but as a mere response to an increase in anxiety.

Our observation indicating the increased gain of sensorimotor feedback control of movements under pressure fits well with the explicit monitoring theory. The theory proposes that after an appraisal of performance pressure, attention shifts exaggeratedly toward the procedural process of motor performance, which eventually collapses skillful performance. Previous studies reported that this attentional shift to the automatized skill execution process is detrimental, particularly for skilled individuals who can perform the well-learned procedural skill without explicitly attending to its underlying process (5, 11). Although this theory explains well that skilled players are better at being aware of a course of skill execution in the higher stressful situation (5), the present study provides another but not exclusive viewpoint that a shift in sensorimotor control from feedforward to feedback control mediates the deleterious effect of pressure on performance. This idea is compatible with a recent finding of a pressure-induced increase in the activation of the anterior cingulate cortex that is responsible for monitoring motor performance (4).

One may speculate that the present observation can be explained by a change in the Kalman gain under pressure in the optimal feedback control (40, 41). In this theory, the Kalman gain in sensorimotor control is modulated based on uncertainty of the sensory feedback (i.e. state estimation). Pianists can accurately predict the sensory consequences of the movements by means of the forward model (42). It is possible that the pressure causes an erroneous state estimation and thereby increases in the Kalman gain, which needs to be investigated in a further experiment that manipulates the uncertainty of the feedback.

Our intervention study clearly demonstrated that practicing the piano while ignoring delayed tone production resulted in robust tempo control against artificially delayed tone production in piano playing under pressure. Similarly, robust sensorimotor control was observed when playing without explicit psychological stress. Therefore, the abnormal elevation of the sensorimotor feedback gain under pressure can be normalized through practicing in this way. The lack of a significant difference in the tempo error between the group that compensated for the delayed tone production and the control group further indicates a specific effect of practicing while ignoring the delayed tone production on the performance fluctuation under pressure. This implicates the potential of the practice of ignoring the auditory perturbation as a training method for stabilizing tempo control in piano performance under pressure. Previous studies demonstrated that skilled singers are better at ignoring the auditory perturbation in singing than nonmusicians (25), similar to pianists (27). The present sensorimotor training may enable pianists to maintain low feedback gain in an uncertain and stressful situation.

There are several possible limitations of the present study. First, because a repetition of playing under pressure potentially causes habituation to psychological pressure, we did not additionally evaluate piano performance without the provision of delayed tone production in the stressful condition. However, this limits the understanding of a direct relationship of the loss of robustness in sensorimotor control with temporal inaccuracy of piano performance in stressful situations, which needs to be further explored. Second, although we evaluated the time interval perception at rest, we did not evaluate time perception while playing the piano, which might be affected by psychological stress, unlike the time perception at rest. Finally, we did not examine changes in finger muscular co-contraction under pressure through ignoring practice, which should be assessed in a future study.

